# Predicting double-strand DNA breaks using epigenome marks or DNA at kilobase resolution

**DOI:** 10.1101/149039

**Authors:** Raphaël Mourad, Olivier Cuvier

## Abstract

Double-strand breaks (DSBs) result from the attack of both DNA strands by multiple sources, including exposure to ionizing radiation or reactive oxygen species. DSBs can cause abnormal chromosomal rearrangements which are linked to cancer development, and hence represent an important issue. Recent techniques allow the genome-wide mapping of DSBs at high resolution, enabling the comprehensive study of DSB origin. However these techniques are costly and challenging. Hence we devised a computational approach to predict DSBs using the epigenomic and chromatin context, for which public data are available from the ENCODE project. We achieved excellent prediction accuracy (*AUC* = 0.97) at high resolution (< 1 kb), and showed that only chromatin accessibility and H3K4me1 mark were sufficient for highly accurate prediction (*AUC* = 0.95). We also demonstrated the better sensitivity of DSB predictions compared to BLESS experiments. We identified chromatin accessibility, activity and long-range contacts as best predictors. In addition, our work represents the first step toward unveiling the”*cis*-DNA repairing” code underlying DSBs, paving the way for future studies of *cis*-elements involved in DNA damage and repair.

## 1 Introduction

Double-strand breaks (DSBs) arise when both DNA strands of the double helix are severed. DSBs are caused by the attack of deoxyribose and DNA bases by reactive oxygen species and other electrophilic molecules [22]. DSBs are particularly hazardous to the cell because they can lead to deletions, translocations, and fusions in the DNA, collectively referred as chromosomal rearrangements [23]. DSBs are most commonly found in cancer cells. Several high-throughput sequencing techniques have been developped for the genome-wide mapping of DSBs *in situ* such as GUIDE-seq [34], BLESS [8] and DSBCapture [17]. The most recent technique, DSBCapture, allowed to map more than 80 thousand endogenous DSBs at a lower than 1 kb resolution in human. To date, DSBs have been mapped at high resolution only for a few number of cell lines, because of high sequencing costs and experimental difficulties. This prevented the comprehensive study of the double-strand break landscape in the human genome across diverse cell lines and tissues.

Chromatin immunoprecipitation followed by high-throughput DNA sequencing (ChIP-seq) and DNase I hypersensitive site sequencing (DNase-seq) data are publicly available for dozens of cell lines with the ENCODE [31] and Roadmap Epigenomics [7] projects. On the one hand, recent studies have shown that the mapping of regulatory elements such as enhancers or promoters can be accurately predicted using available epigenome and chromatin data [9,14]. Other studies have shown that the epigenome can in its turn be predicted by combinations of DNA motifs and DNA shape [21, 29, 36, 38]. On the other hand, DSBs and the resulting DNA repair mechanisms were shown to be linked to epigenome marks, including H3K4me1/2/3 and chromatin accessibility [17]. Accordingly, PRDM9-mediated trimethylation of H3K4 (H3K4me3) was originally shown to play a critical role in regulating DSBs associated with meiotic recombination hotspots [1, 11, 24]. Moreover the repair of DSBs involves both posttranslational modification of histones, in particular γ-H2AX, and concentration of DNA-repair proteins at the site of damage [13, 28]. It remains unclear to what extent DNA motifs or histone modifications predict or regulate the cellular response to DSBs in other developmental stages. Here we thus sought to test whether publicly available epigenome and chromatin data, or DNA motifs and shape could be used to possibly predict DSBs.

In this article, we demonstrate, for the first time, that endogenous DSBs can be computationally predicted using the epigenomic and chromatin context, or using DNA sequence and DNA shape. Our predictions achieve excellent accuracy (*AUC* > 0.97) at high resolution (<1kb) using available ChIPseq and DNase-seq data from public databases. DNase, CTCF binding and H3K4me1/2/3 are among the best predictors of DSBs, reflecting the importances of chromatin accessibility, activity and long-range contacts in determining DSB sites and subsequent repairing. Another important predictor is p63 binding, a member of the p53 gene family known to be involved in DNA repair. We also successfully predict DSB sites using DNA motif occurences only (*AUC* = 0.839), supporting a”cis-DNA repairing” code of DSBs involving numerous DNA damage and repair protein binding motifs including members of the transcription factor complex AP-1 and p53 family. In addition, DNA shape analysis further reveals the importance of the structure-based readout in determining DSB sites, complementary to the sequence-based readout (motifs).

## 2 Results and Discussion

### 2.1 Double-strand break prediction approach

Our computational approach to predict DSBs is schematically illustrated in Figure 1. In the first step, we analyzed public DSBCapture data from Lensing *et al*. [17], which provided the most sensitive and accurate genome-wide mapping of double-strand breaks to date (Figure 1a). DSBCapture is a technique that captures DSBs *in situ* and that can directly map them at the single-nucleotide resolution. DSBCapture peaks were called with less than 1 kb resolution (median size of 391 bases). The DSBCapture peaks obtained from two biological replicates were intersected to yield more reliable DSB sites. Endogeneous breaks were captured for NHEK cells, for which numerous ChIP-seq and DNase-seq data were made publicly available by the ENCODE project [31]. In the second step, we integrated and mapped different types of data within DSB sites and non-DSB sites. To prevent bias effects, non-DSB sites were randomly drawn from the human genome with sizes, GC and repeat contents similar to those of DSB sites [10] (Figure 1b). ChIP-seq and DNase-seq peaks in NHEK cells as obtained from the ENCODE project were mapped corresponding to DSB and non-DSB sites [31]. We also mapped p63 ChIP-seq peaks from keratinocyte cells (HKC) [15]. We further searched for potential protein binding sites at DSB and non-DSB sites using JASPAR 2016 database motif position weight matrices [20], and predicted DNA shape at DSB and non-DSB sites using Monte Carlo simulations [6]. In the third step, a random forest classifier was built to discriminate between DSB sites and non-DSB sites based on epigenome marks or DNA (Figure 1c). Random forest variable importances were used to estimate the predictive importance of a feature. We also compared random forest predictions with another popular method, lasso logistic regression [32]. Using lasso regression, we assessed the positive, negative or null contribution of a feature to DSBs. We then split the DSB dataset into a training set to learn model parameters by cross-validation, and into a testing set to compute receiver operating characteristic (ROC) curve and the area under the curve (AUC) for prediction accuracy evaluation.

**Figure 1.**
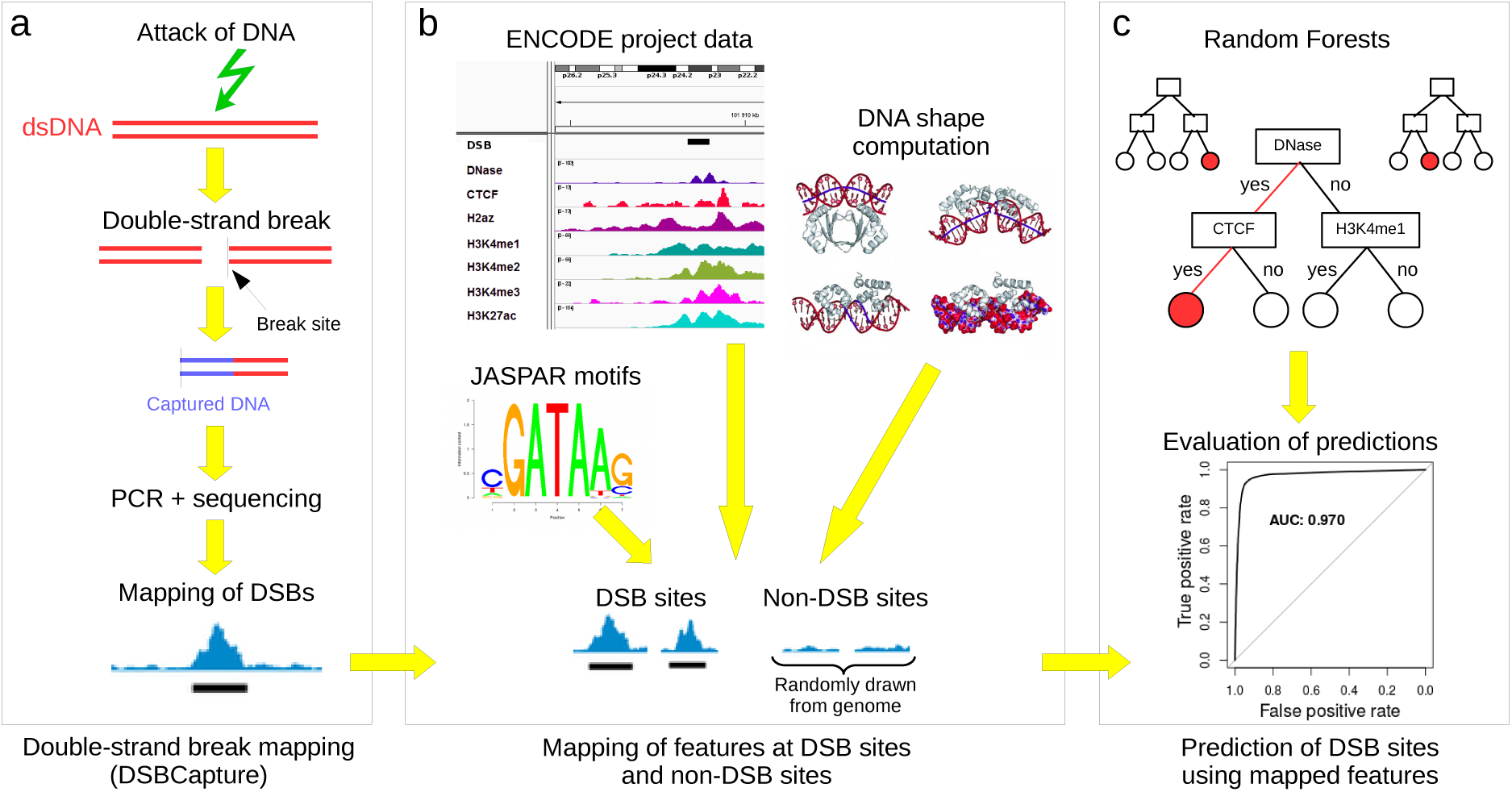
Double-strand break (DSB) prediction using epigenome mark or DNA. The prediction approach consisted in three steps. a) Mapping of DSBCapture sequencing data and DSB peak calling. b) Mapping of features at DSB and non-DSB sites. Features included epigenomic and chromatin data from ENCODE project, DNA motifs from JASPAR database and DNA shape predictions. c) Prediction of DSB sites using features.

### 2.2 Double-strand breaks are enriched with epigenome marks and DNA motifs

We first sought to comprehensively assess the link between DSBs and epigenome marks or DNA motifs. As previously shown [17,30], several epigenomic and chromatin marks colocalized at double-strand breaks (Figure 2a). Among the most enriched marks were DNase I hypersensitive sites, H3H4 methylations and CTCF (Figure 2b). For instance, 91% of DSBs colocalized to a DNase site, whereas this percentage dropped to 11% for non-DSB regions. This corresponded to an odds ratio (*OR*) of 89.3. Similarly a high enrichment was found for H3K4me2 (74% versus 11%; *OR* = 22.4) and for the insulator protein CTCF (25% versus 2%; *OR* = 19), which may involve its interactions with the insulator-related cofactor cohesin that has been shown to protect genes from DSBs [5]. As such, DSBs mostly localized within open and active regions that were often implicated in long-range contacts [27]. Interestingly, DSBs also colocalized with tumor protein p63 binding (19.4% versus 1%; OR = 23.8), a member of the p53 gene family involved in DNA repair [19,37]. In addition, we could distinguish DNase and CTCF sites that were enriched at the center of DSBs, from histone marks that were found at the edges of DSB sites (Figure 2c). Therefore the strong enrichments of epigenomic and chromatin marks at DSB sites suggested that DSB regions could be accurately predicted using available ChIP-seq and DNase-seq data from public databases including ENCODE and Roadmap Epigenomic.

**Figure 2.**
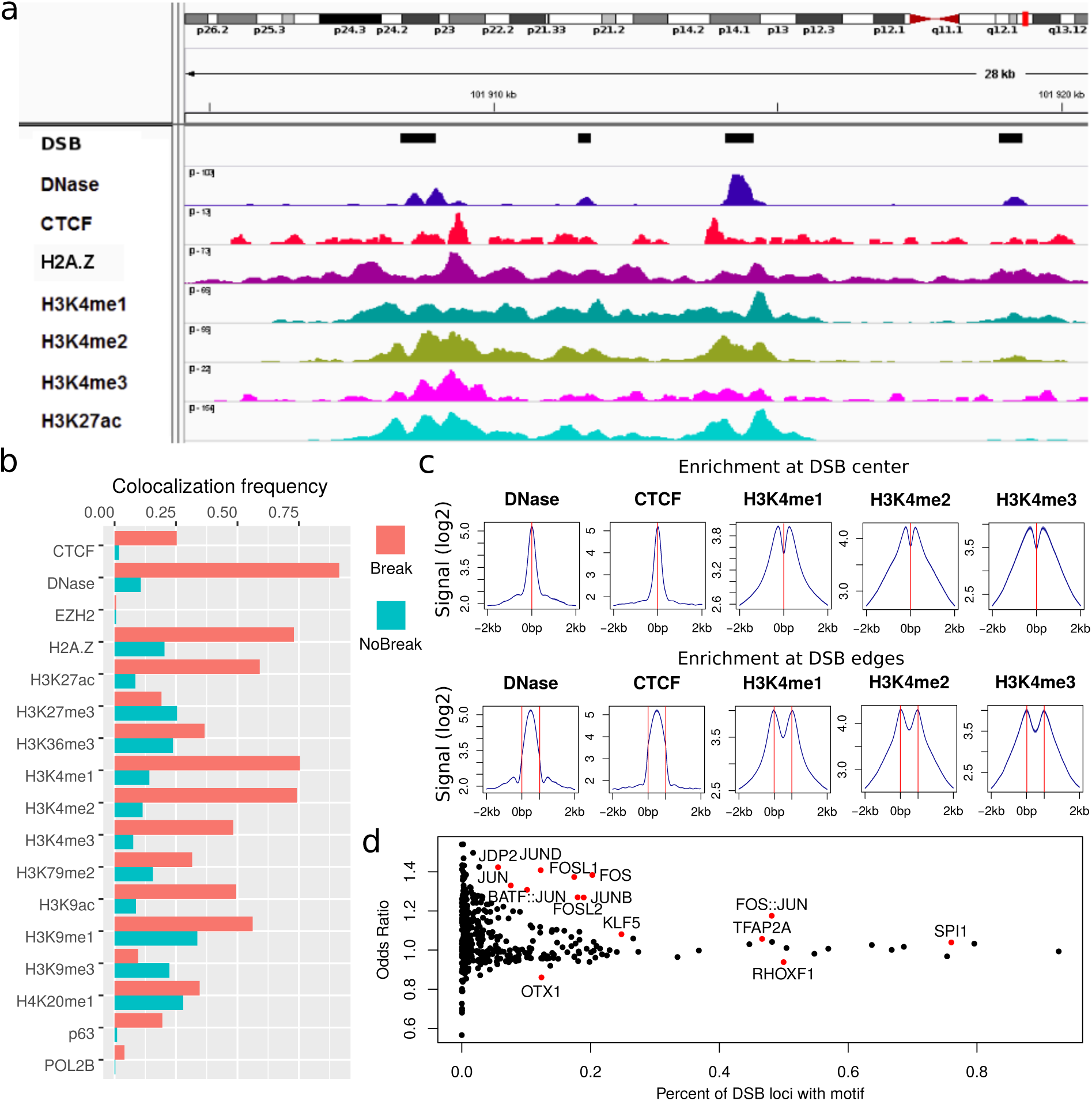
Epigenomic, chromatin and DNA motif profiles of double-strand breaks (DSBs). a) A genome browser view of DSBs with histone marks, chromatin openess (DNase-seq) and DNA binding proteins. b) Colocalization frequencies of epigenomic marks and DNA binding proteins at DSB sites, as compared to non-DSB sites. c) Average profiles of epigenomic marks and DNA binding proteins at DSB sites. d) Enrichment of DNA motifs at DSB sites, as measured by the odds ratio and the percent of DSB loci with motif.

Previous enrichment analyses of DNA-binding proteins were limited by available ChIP-seq data. Hence we sought which DNA motifs may be enriched at DSB sites as a way to obtain a more comprehensive list of candidate DNA-binding proteins. Over the 434 available motifs from JASPAR database, 134 were significantly enriched (*p <* 0.05, Bonferroni correction), indicating that DSBs were associated with a large number of protein binding sites (Figure 2d). Among the most enriched and frequent motifs, we identified numerous motifs specifically recognized by protein cofactors of the transcription factor complex AP-1 whose activity has been shown to be induced by genotoxic agents. This included JUND (*OR* = 1.40, 12% of DSBs), JUNB (*OR* = 1.27, 19% of DSBs), the heterodimer BATF::JUN (*OR* = 1.31, 10% of DSBs), and also FOS (*OR* = 1.37, 20% of DSBs), FOSL1 (*OR* = 1.37, 17% of DSBs) and FOSL2 (*OR* = 1.27, 18% of DSBs). Among the most enriched but less frequent motifs, we found as expected CTCF (*OR* = 1.54, 1.7% of DSBs), as well as the members of the tumor protein family p53, *i.e*. p53 itself (*OR* = 1.54, 0.2% of DSBs), p63 (*OR* = 1.49, 0.3% of DSBs) and p73 (*OR* = 1.54, 0.1% of DSBs), whose cofactors are specifically involved in the response to DNA damage [19, 37]. Such enrichments of DNA motifs at DSB sites therefore supported the view that DNA sequence could already predict some of the DSBs encountered.

### 2.3 Prediction using epigenomic and chromatin data

Given the strong link between DSBs and epigenomic and chromatin marks, we sought to build a classifier to discriminate between DSB sites from non-DSB sites based on the presence/absence of such marks. For this purpose, we used random forests that represent very efficient classifiers to predict a feature, and that can capture non-linear and complex interaction effects [3]. Using this classifier, we obtained excellent predictions of DSBs based on the epigenomic and chromatin marks available (AUC=0.970; Figure 3a). We could also compute the “variable importance” (*VI*) that reflects the importance of a mark as a predictor (Figure 3b). Among the marks, DNase showed the highest variable importance (*VI* = 0.180), reflecting known higher chromatin accessibility after DNA damage [28] or the involvement of chromatin remodeling complexes in processing of DSBs [12]. Other good predictors were CTCF (*VI* = 0.042), p63 (*VI* = 0.031), H3K4me1 (*VI* = 0.028), H3K4me2 (*VI* = 0.019), H3K4me3 (*VI* = 0.012) and H3K27ac (*VI* = 0.010), highlighting the roles of active chromatin, but also long-range contacts and DNA damage response in predicting DSB sites. A drawback of variable importance lies in its inability to distinguish between the positive or negative contribution of the predictive mark on DSBs. For this reason, we also used lasso logistic regression to predict DSBs [32]. With this second model, we obtained excellent predictions, although slightly less accurate (AUC=0.967, Supplemental Figure 1). From lasso regression, we could assess the positive or negative contributions of the predictive marks using beta coefficients (Figure 3c). We identified positive predictive contributions of DNase, CTCF, p63, H3K4me1 and H3K4me2 marks, as previously revealed by enrichment analysis. We also uncovered negative predictive contributions of H3K9ac, H3K36me3 and H3K79me2. In agreement, H3K9ac was shown to be rapidly and reversibly reduced in response to DNA damage [33]. Moreover, H3K36me3 may negatively impede on DSBs by restricting chromatin accessibility through nucleosome positioning [18] or more directly by favoring the repair of DSBs [26].

**Figure 3.**
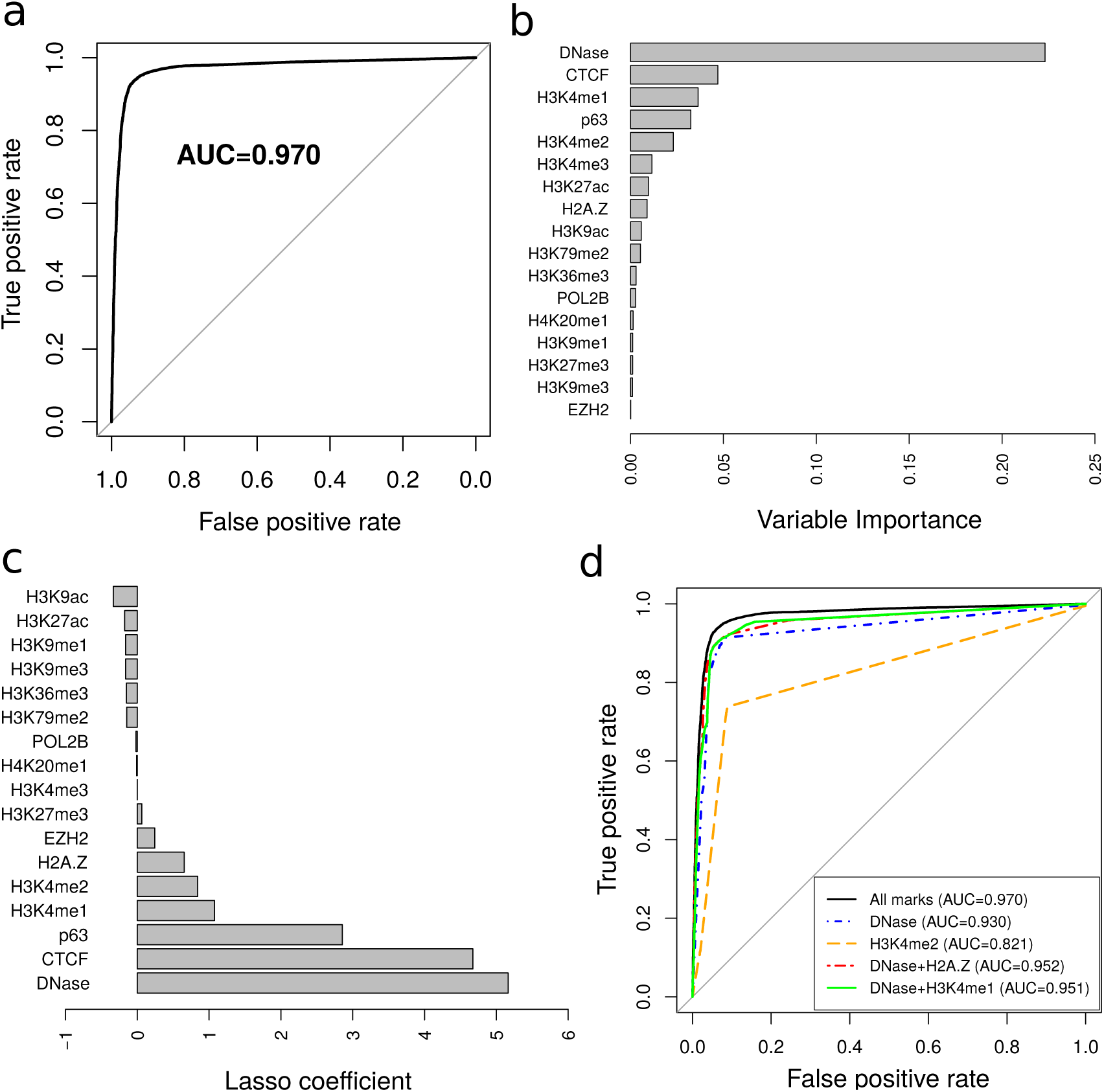
Prediction of double-strand breaks (DSBs) using epigenomic marks and random forests. a) Receiver operating characteristic (ROC) curve for the prediction of DSBs. Area under the curve (AUC) is plotted. b) Variable importances of epigenomic marks. c) Lasso logistic regression coefficients. d) Different predictive models including all marks, DNase only, H3K4me2 only, DNase+H2A.Z, or DNase+H3K4me1.

We next sought to build a classifier using only one or two epigenomic marks, because it could allow to predict DSB sites even for cells for which only a few data may be available. We found that DNase I sites alone were sufficient to achieve good prediction accuracy (AUC=0.919), whereas H3K4me2 was not sufficient (AUC=0.816). Combinations of DNase with H2A.Z or H3K4me1 yielded very accurate predictions (AUC=0.952 and AUC=0.951, resp.), close to the model including all marks. All results demonstrated that DSBs can be accurately predicted at less than 1 kb resolution using just a few data.

### 2.4 Model predictions outperformed BLESS experiment and were validated using independent dataset

We then compared previous DSB predictions with DSBs identified by BLESS experiments [8,17]. We also included in the comparison DSBCapture DSBs as gold standard. We first looked at predicted DSB sites surrounding the two genes MYC and MAP2K9 (Figure 4a). For MYC, random forests correctly identified the 4 DSBs that were detected by DSBCapture, but erroneously predicted one DSB (yellow circle), whereas BLESS only identified one DSB out of four. For MAP2K3, random forests successfully predicted all DSBs detected by DSBCapture, whereas BLESS only identified three DSBs out of 11. We then compared predictions with BLESS at the genome-wide level (Figure 4b). We observed that random forests correctly predicted 18084 out of 18510 DSB sites (97.70%) found by BLESS, while it also successfully identified additional 63587 out of 66593 DSB sites (95.48%) found by DSBCapture that were not detected by BLESS. The model only misclassified 1552 sites as DSBs. Such comparisons thus revealed the better sensitivity of random forest predictions in detecting DSBs compared to BLESS experiments.

**Figure 4.**
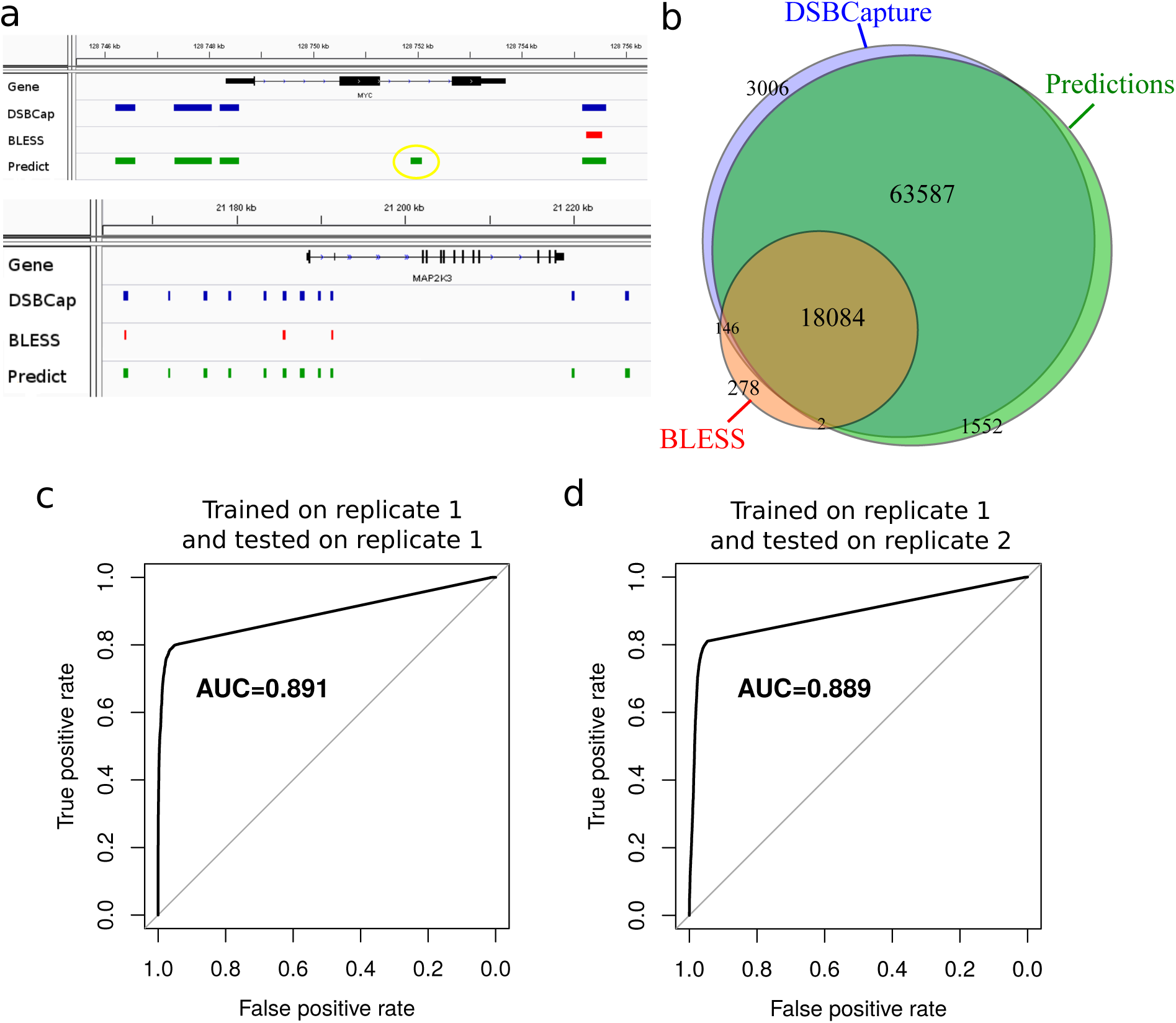
Comparison of predicted and BLESS double-strand breaks (DSBs) and validation with an independent dataset. a) Comparison for the MYC and MAP2K9 genes. b) Venn diagram illustrating the overlaps between DSBCapture (gold standard), random forest predictions and BLESS DSBs. c) Receiver operating characteristic (ROC) curve for the prediction of DSBs trained on replicate 1 and tested on same replicate. Area under the curve (AUC) is plotted. d) ROC curve for the prediction of DSBs trained on replicate 1 and tested on replicate 2.

In the previous subsection, we evaluated the accuracy of model predictions using a testing dataset that was from the same data as the training data (DSBs that overlapped between two replicates were split into a training and a testing datasets). Here we assessed model predictions by training random forests on a first biological replicate and by testing prediction accuracy on a second biological replicate. For this purpose, we used the two available DSBCapture biological replicates [17]. Accordingly, we used ENCODE epigenomics data for which two biological replicates were available: DNase, CTCF, H3K4me3, H3K27me3 and H3K36me3. The first (resp. second) replicates of ENCODE data were associated with the first (resp. second) DSBCapture replicate. Using only those 5 DNase-seq and ChIP-seq data, the model learned with the first replicate achieved accurate predictions on tested data from the first replicate (AUC=0.891) (Figure 4c). It is noteworthy that the observed lower accuracy compared to previous subsection (Figures 3a and d) can be explained by the small number of epigenomic data, and the lower reliability of DSBs identified using only one DSBCapture replicate. To validate the model on an independent dataset, we predicted DSBs from the second replicate using the model trained on the first replicate together with DNase-seq and ChIP-seq data of the second replicate. We obtained accurate predictions (AUC=0.889) close to the one obtained for the first replicate (Figure 4d). These accurate predictions demonstrated that using a classifier trained with epigenome and chromatin data represented a reliable strategy for predicting DSBs.

### 2.5 Prediction from DNA motifs and shape

There is growing evidence supporting the importance of the *cis*-regulatory code in shaping the epigenome [36]. Moreover, we previously identified a large number of DNA motifs that were enriched or depleted at DSB sites, suggesting a “*cis*-DNA repairing” code. Hence, we explored the possibility to predict DSBs based on the occurrence of DNA motifs from JASPAR database. We built a random forest classifier using 434 available motifs and obtained good prediction accuracy (AUC=0.827; Figure 5a). Several motifs from the transcription factor complex AP-1 represented good predictors such as FOS::JUN (*VI* = 0.016) and FOS (*VI* = 0.009) (Figure 5b), which were previously shown to be enriched at DSB sites. We also uncovered TFAP2A motif as good predictor (*VI* = 0.011), corresponding to a protein that has not been linked to DNA repair yet. Using lasso regression, we improved previous predictions (AUC=0.839; Figure 5a). Based on lasso regression, we found that the CTCF motif presented the highest beta coefficient (*β* = 3.22), corresponding to an odds ratio *OR* = 25 (Figure 5c), supporting recent evidence showing that long-range contacts are involved in DNA repair [2,30]. Furthermore we found many motifs recognized by proteins that participate to DNA damage and repair. For instance, all tumor proteins p53, p63 and p73 motifs showed high coefficients (*β* > 2.03, *OR* > 7.6). Interestingly, we also found motifs recognized by factors highly related to DSB pathways such as those involved in heavy metal response (MTF-1: *β* = 2.08, *OR* = 8), in oxidative stress response (NRF1: *β* = 0.93, *OR* = 2.53; REST: *β* = 1.75, *OR* = 5.75), in endoplasmic reticulum stress (ATF4: *β* = 0.97, *OR* = 2.64) and in estrogen-induced DNA damage (ESR1: *β* = 0.88, *OR* = 2.41). Many of the abovementionned proteins were actually shown to interact with each other. For instance, NRF1 associates with Jun proteins of AP-1 complex [35]. ESR1 associates with AP-1/JUN and FOS to mediate estrogen element response (ERE)-independent signaling [16], which may thus participate to coordinate multiple functions along with the processing of DSBs.

DNA shape was recently shown to predict transcription factor binding sites and gene expression [21, 25]. We thus assessed if DNA shape could similarly serve to predict DSBs together with motifs. For this purpose, we predicted four DNA shape features using simulations: minor groove width (MGW), propeller twist (ProT), roll (Roll) and helix twist (HelT) of DSB sites at base resolution. From each feature, we computed 12 predictors including quantiles (0%, 10%, 20%, 30%, 40%, 50%, 60%, 70%, 80%, 90% and 100%) and the variance to describe the distribution of the feature within a DSB site. We used the resulting 48 variables combined with motif occurences to predict DSBs with random forests and obtained better accurary (*AUC* = 0.838) compared to using motifs alone (*AUC* = 0.827; Figure 5d). Among the DNA shape variables, ProT median and MGW variance presented the highest variable importances (*VI* = 0.01 and *VI* = 0.01, resp.). Using lasso regression, we also obtained better predictions (*AUC* = 0.858), compared to using motifs only (*AUC* = 0.839) (Figure 5d). These results reflected the importance of DNA shape in determining DSB sites, in agreement with studies showing that narrow minor grooves (created by either sequence context or DNA bending) limit access by reactive oxygen species [4].

**Figure 5.**
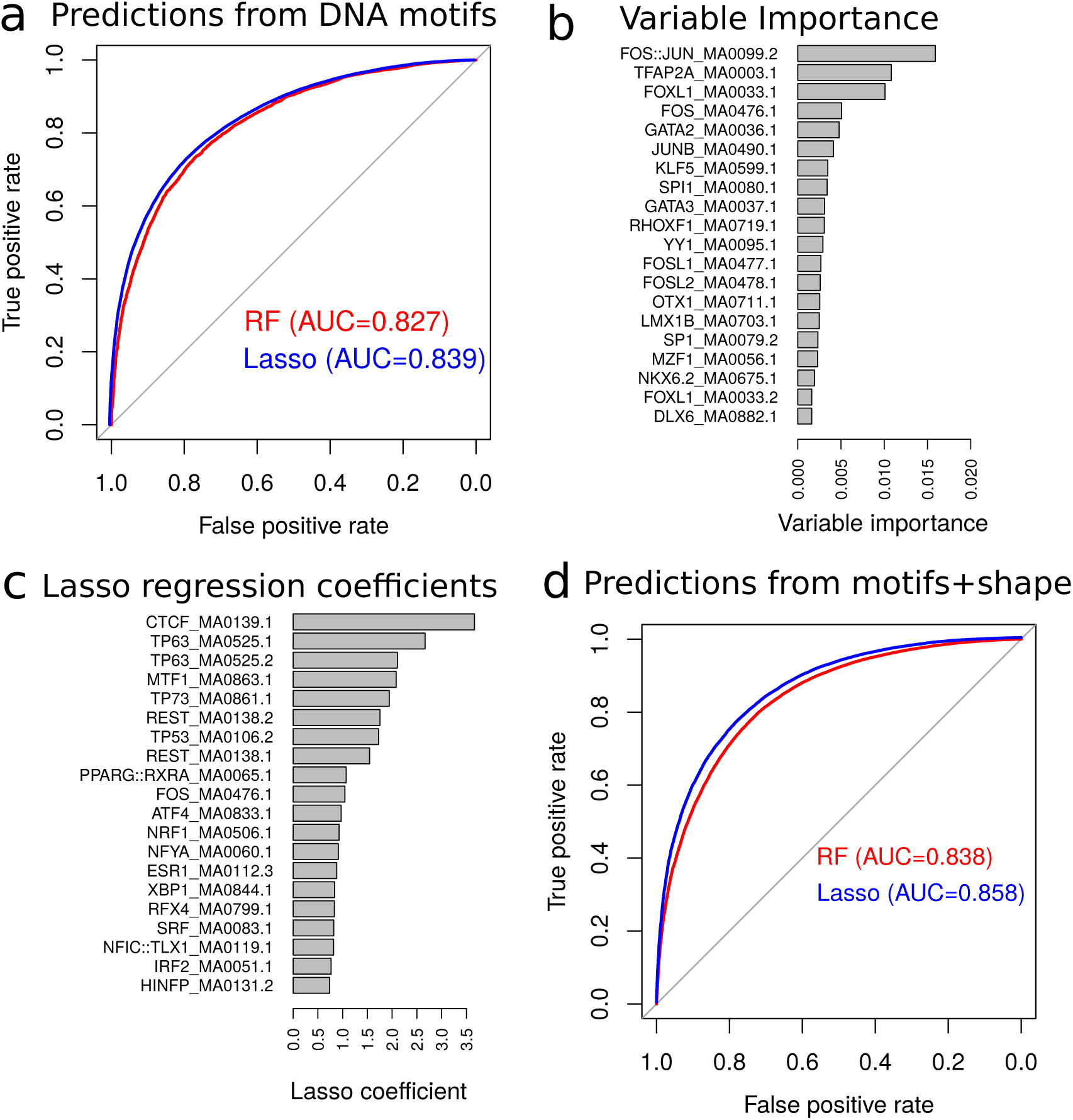
Prediction of double-strand breaks (DSBs) using DNA motifs and shape. a) Receiver operating characteristic (ROC) curve for the DSB predictions using DNA motifs from JASPAR database. Random forest (RF) and lasso logistic regression (Lasso) were compared. b) The 20 highest DNA motif variable importances. c) The 20 highest DNA motif lasso coefficients. d) ROC curve for the DSB predictions using DNA motifs with DNA shape.

## 3 Conclusion

Double-strand breaks represent a major threat to the cell, and they are associated with cancer development. Here we show, for the first time, that such DSBs can be computationally predicted using public epigenomic data, even when the availability of data is limited (e.g. DNase I and H3K4me1). By using state-of-the-art computational models, we achieve excellent prediction accuracy, paving the way for a better understanding of DSB formation depending on developmental stage or cell-type specific epigenetic marks. In addition, our work represents the first step toward unveiling the “*cis*-DNA repairing” code underlying DSBs, which is composed of numerous DNA motifs for binding of key regulators of DNA repair, and could guide further locus-specific genome editing.

## 4 Materials and Methods

### 4.1 Double-strand breaks

We used double-strand breaks mapped by DSBCapture in human epidermal keratinocytes (NHEK) cells [17]. DSBCapture peaks were called using MACS 2.1.0 on human genome assembly hg19 (https://github.com- /taoliu/MACS). The peaks obtained from two biological replicates were intersected to yield more reliable DSB sites used for model predictions.

### 4.2 ChIP-seq and DNase-seq data

We used ChIP-seq (CTCF, POL2B, EZH2, H3K4me1/me2/me3, H3K9me1/me3/ac, H3K27me3/ac, H3K36me3, H4K20me1) and DNase-seq data for NHEK cells from the ENCODE project [31]. We also used p63 ChIP-seq of keratinocyte cells (HKC) from Kouwenhoven *et al*. [15].

### 4.3 DNA motifs

We used transcription factor binding site (TFBS) motif position frequency matrices from the JAS-PAR database (http://jaspar.genereg.net/). We called transcription factor binding sites over the human genome using the position weight matrices and a minimum matching score of 80%.

### 4.4 DNA shape

We predicted four DNA shape features using Monte Carlo simulations: minor groove width (MGW) and propeller twist (ProT) at base pair (bp) resolution and values of roll (Roll) and helix twist (HelT) at base pair step resolution using R package DNAshapeR (https://bioconductor.org/packages/release/bioc/html /DNAshapeR.html).

### 4.5 Random forest and lasso regression

We used R package ranger (https://cran.r-project.org/web/packages/ranger/) to efficiently compute random forest classification [3]. We used the default package parameters: *num.trees* = 500 and *mtry* is the square root of the number variables. Variable importance was computed using the mean decrease in accuracy in the out-of-bag sample. To discrimate between DSB and non-DSB sites, we randomly selected genomic sequences that matched sizes, GC and repeat contents of DSB sites using R package gkmSVM (https://cran.r-project.org/web/packages/gkmSVM/). To learn the model, we mapped epigenomic data, DNA motifs and DNA shape as follows. For epigenomic data including ChIP-seq and DNase-seq data, we used peak genomic coordinates of a feature (for instance CTCF binding sites) and considered the presence (*x* = 1) or absence (*x* = 0) of the corresponding feature at the DSB site. If a feature peak only overlapped 60% of the DSB site, then *x* = 0.6. For DNA motifs, we computed the number of motif occurrence within DSB and non-DSB sites. For DNA shape, we computed 4 features including MGW, ProT, Roll and HelT of DSB sites at base resolution. For each DNA shape feature, we then computed 12 predictors including quantiles (0%, 10%, 20%, 30%, 40%, 50%, 60%, 70%, 80%, 90% and 100%) and the variance to describe the distribution of the feature within a DSB site. The DSB data were next split into two sets: the training set used for learning the model and a test set used for assessing prediction accuracy. We also used R package glmnet (https://cran.r-project.org/web/packages/glmnet/index.html) to compute lasso logistic regression with cross-validation. To estimate prediction accuracy of random forests and lasso regression, we computed the receiver operating characteristic (ROC) curve and the area under the curve (AUC).

## Acknowledgements

The authors are grateful to the Balasubramanian lab (Babraham Institute, UK) for data availability and for help in processing them. The authors are also grateful to the genotoul bioinformatics platform Toulouse Midi-Pyrenees for providing computing resources.

## 5 Supplemental Files

**Supplemental Figure 1.**
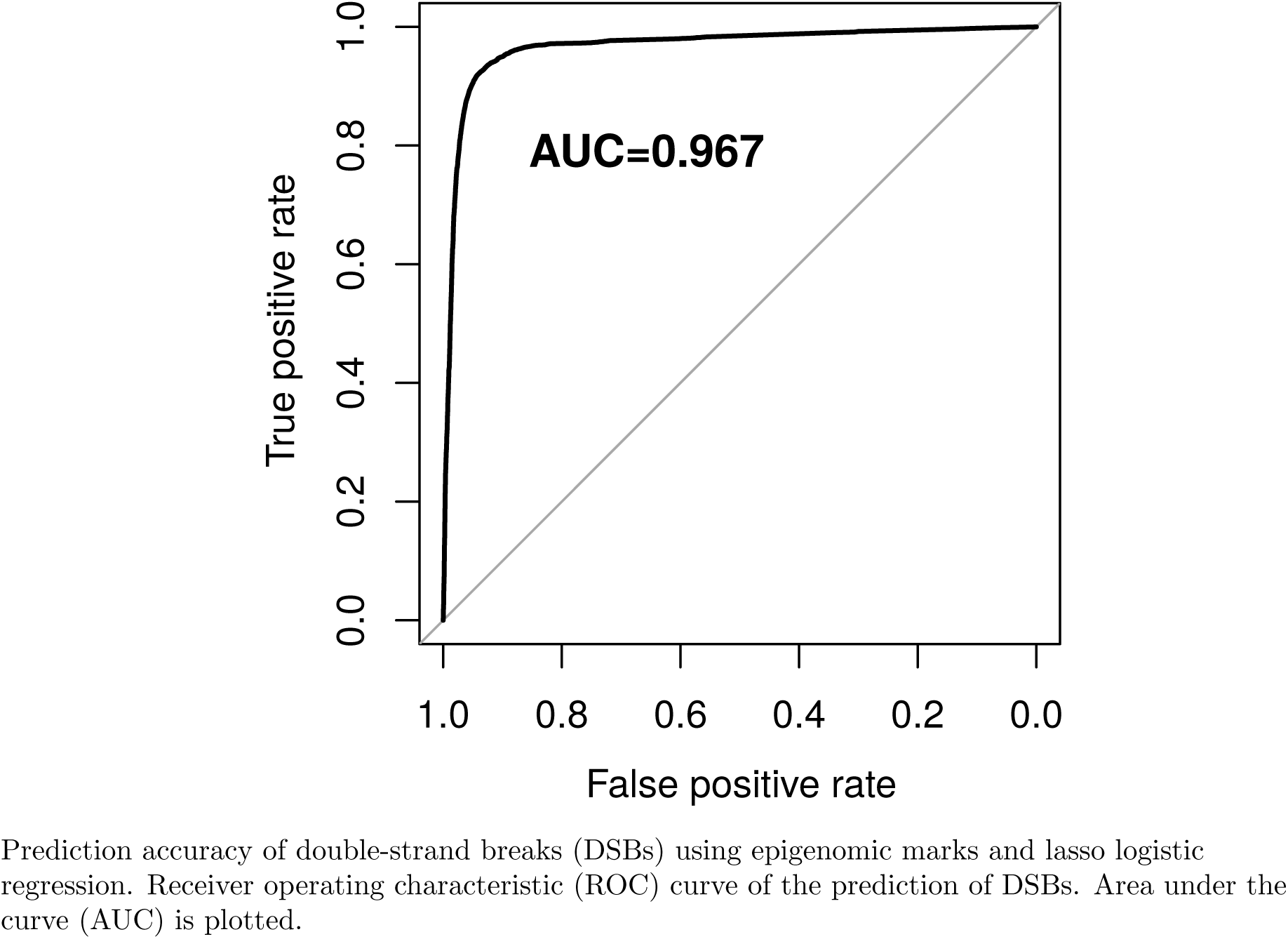
Prediction accuracy of double-strand breaks (DSBs) using epigenomic marks and lasso logistic regression. Receiver operating characteristic (ROC) curve of the prediction of DSBs. Area under the curve (AUC) is plotted.

